# A humanized knock-in *Col6a1* mouse recapitulates a deep-intronic splice-activating variant

**DOI:** 10.1101/2024.03.21.581572

**Authors:** Véronique Bolduc, Fady Guirguis, Berit Lubben, Lindsey Trank, Sarah Silverstein, Astrid Brull, Matthew Nalls, Jun Cheng, Lisa Garrett, Carsten G. Bönnemann

## Abstract

Antisense therapeutics such as splice-modulating antisense oligonucleotides (ASOs) are promising tools to treat diseases caused by splice-altering intronic variants. However, their testing in animal models is hampered by the generally poor sequence conservation of the intervening sequences between human and other species. Here we aimed to model in the mouse a recurrent, deep-intronic, splice-activating, *COL6A1* variant, associated with a severe form of Collagen VI-related muscular dystrophies (COL6-RDs), for the purpose of testing human-ready antisense therapeutics *in vivo*. The variant, c.930+189C>T, creates a donor splice site and inserts a 72-nt-long pseudoexon, which, when translated, acts in a dominant-negative manner, but which can be skipped with ASOs. We created a unique humanized mouse allele (designated as “h”), in which a 1.9 kb of the mouse genomic region encoding the amino-terminus (N-) of the triple helical (TH) domain of collagen α1(VI) was swapped for the human orthologous sequence. In addition, we also created an allele that carries the c.930+189C>T variant on the same humanized knock-in sequence (designated as “h+189T”). We show that in both models, the human exons are spliced seamlessly with the mouse exons to generate a chimeric mouse-human collagen α1(VI) protein. In homozygous *Col6a1* ^h+189T/h+189T^ mice, the pseudoexon is expressed at levels comparable to those observed in heterozygous patients’ muscle biopsies. While *Col6a1*^h/h^ mice do not show any phenotype compared to wild-type animals, *Col6a1* ^h/h+189T^ and *Col6a1* ^h+189T/h+189T^ mice have smaller muscle masses and display grip strength deficits detectable as early as 4 weeks of age. The pathogenic h+189T humanized knock-in mouse allele thus recapitulates the pathogenic splicing defects seen in patients’ biopsies and allows testing of human-ready precision antisense therapeutics aimed at skipping the pseudoexon. Given that the *COL6A1* N-TH region is a hot-spot for COL6-RD variants, the humanized knock-in mouse model can be utilized as a template to introduce other *COL6A1* pathogenic variants. This unique humanized mouse model thus represents a valuable tool for the development of antisense therapeutics for COL6-RDs.

## Introduction

Collagen VI-related muscular dystrophies (COL6-RDs) form a group of disorders that span a broad severity spectrum ranging from the debilitating and severe Ullrich type to the milder Bethlem form, connected by intermediate phenotypes. Characteristic manifestations include progressive muscle weakness, worsening proximal joints contractures, distal joints hyperlaxity, and progressive respiratory failure. In Ullrich type, the disease manifests at birth and without ventilatory support, can be fatal in the first two decades of life. In the milder cases, symptoms can become evident in young adulthood with slower progression but with no impact on life expectancy (reviewed in (1, 2)). Symptoms are currently managed with physical therapy and ventilatory support, but there are no specific therapies available yet.

COL6-RDs are caused by the absence or dysfunction of collagen VI, a microfibrillar extracellular matrix protein abundant in several organs including skeletal muscles where it surrounds myofibers (1). The three main genes, *COL6A1, COL6A2,* and *COL6A3*, encode the three major collagen VI alpha peptide chains (α1(VI), α2(VI), and α3(VI)) (1, 3) and are mainly expressed by the interstitial fibro-adipogenic progenitor cells (FAPs) (4–6). In FAPs, the three chains intertwine through their respective TH domains, proceeding from their C-towards their N-termini, to form monomers. The monomers form dimers and tetramers, which are secreted into the extracellular space where they polymerize and form the collagen VI meshwork (reviewed in (7, 8)).

Pathogenic *COL6* variants can be inherited in an autosomal recessive manner, but, more frequently, they are acquired *de novo* and act dominant negatively (1, 7). The most frequent dominant-negative variants cause either glycine substitutions in the Gly-X-Y repeats or in-frame exon deletions, and they occur predominantly in the exons encoding the N-termini TH (N-TH) of any of the three α(VI) peptides. In this location, they allow the mutant collagen α(VI) chains to incorporate into monomers, and further dimers and tetramers, but they affect the function of the collagen VI tetramers in the matrix, hence their dominant-negative effect (1, 7). More recently, a surprisingly common and recurrent pathogenic variant has been identified in intron 11 of *COL6A1*: c.930+189C>T. This intronic variant is also located within the N-TH-encoding region and creates a splice donor (SD) site that, combined with an upstream cryptic splice acceptor (SA) site, inserts an in-frame 72-nt-long pseudoexon into the mature mRNA transcript (9). We previously showed that the pseudoexon is only inserted in approximately 50% of the mRNA transcribed from the +189T allele, yet, this expression level is sufficient to have a dominant-negative effect on collagen VI assembly and to cause the severe clinical manifestations of Ullrich type of COL6-RDs (10, 11). COL6-RDs downstream pathophysiology in muscle is complex and not fully understood (7, 8); therefore, addressing the disease upstream at the genetic source, such as with ASOs, is a promising therapeutic approach. We have shown that skipping the pseudoexon with ASOs restores nearly healthy collagen VI matrix in cultured patient fibroblasts (10), but *in vivo* testing has not been feasible because of the unavailability of a suitable animal model given that human intronic sequences significantly diverge from the corresponding mouse sequences.

Genomic humanization involves the introduction of human genomic sequences, coding and non-coding regulatory sequences included, into a trans-species genome (12). While the process for obtaining these humanized loci was previously tedious, with the currently available genome editing technologies it can be achieved rapidly (12, 13). Entire – or nearly entire – human genes have thus been successfully introduced, and expressed, either in a transgenic location (14), or at the endogenous locus in the trans-species genome, substituting their orthologous counterpart (15). Partial genomic humanization has also been successfully applied (16). Some of these models have proven invaluable to study basic gene regulation and function in the context of human health and disease (12). More recently, humanized animals have been proposed as models to introduce human pathogenic variants and test precision medicine therapeutics (17, 18).

Here we report the partial genomic humanization of the N-TH domain of the mouse *Col6a1* gene (encompassing 1.9-kb and spanning from intron 8 to intron 14). This region includes the relevant exonic and intronic sequences for dominantly acting variants in *COL6A1*, including the pseudoexon-inducing c.930+189C>T variant. We created a humanized wild-type allele (harboring the reference sequence), and a humanized pathogenic allele (carrying the c.930+189T variant), and validated their use as preclinical models with molecular, biochemical, histological, and behavioral assays.

## Results

### Generation of two genomic humanized alleles in *Col6a1*

The variant to model (*COL6A1* c.930+189C>T) is schematized in Figure 1A. *COL6A1*/*Col6a1* coding sequences are highly conserved between human and mouse with 83.2% nucleotide (NM_001848.3 and NM_009933.5) and 90% amino acid (NP_001839.2 and NP_034063.1) sequence identities. In contrast, the intronic sequences are poorly conserved. For instance, the alignment of the human intron 11 sequence with the mouse sequence shows 45.1% identity and 44.3% of gaps. Importantly, the ‘aggc’ site that is converted to ‘AGgt’ due to the c.930+189C>T variant is absent in the mouse intron 11 sequence. Thus, we sought to substitute part of the mouse *Col6a1* gene with the human sequence. We replaced a 1,794-nt-long segment of the mouse *Col6a1* gene (encompassing introns 8 to 14) with the 1,934-nt-long orthologous human sequence (Figure 1B). We generated two humanized knock-in alleles, carrying either the reference cytosine at position c.930+189 (called “h”), or the pathogenic thymine (called “h+189T”) (Figure 1B). The humanized protein sequence spans from mouse Gly268 to Asp351 and includes the Cys344, important for collagen VI dimerization.

**Figure 1.**
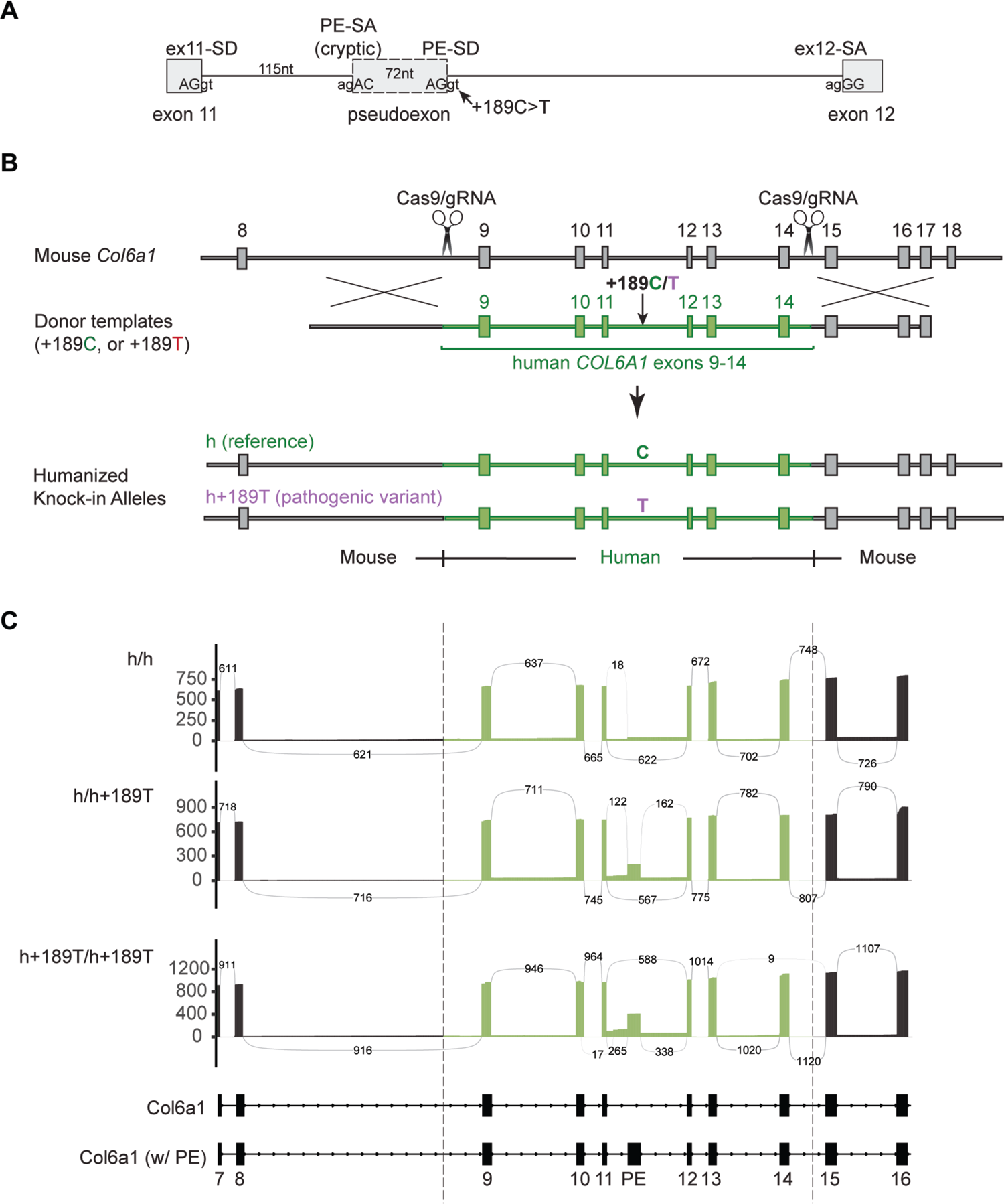
Generation and validation of a *Col6a1* humanized knock-in mouse to model the c.930+189C>T pseudoexon-inducing variant. **(A)** Schematic of the variant to model: the c.930+189C>T variant in intron 11 of the *COL6A1* gene, which creates a donor splice site (PE-SD) and which, upon concomitant activation of an upstream cryptic splice acceptor site (PE-SA), causes the insertion of a 72-nucleotide-long pseudoexon (PE). **(B)** Two *Col6a1* humanized knock-in alleles were generated using a CRISPR/Cas9-induced homologous recombination strategy to swap the mouse genomic sequence spanning the region between introns 8 and 14 with the orthologous human sequence. One allele contains the reference cytosine at position +189 (referred to as allele ‘h’), and the other allele contains the pathogenic thymine (referred to as ‘h+189T’). **(C)** *Col6a1* transcripts were enriched using a probe capture method prior to long-read RNA sequencing. Sashimi plots depict the splicing of human exons with the mouse exons in quadriceps of 8-week-old males.

### Expression and splicing of the *Col6a1* hybrid gene

With long-read RNA sequencing, human exons 9 to 14 were found to seamlessly splice with the mouse exons and to generate full-length mouse-human chimeric *Col6a1* transcripts in quadriceps tissue (Figure 1C, Supplementary Figure S1). Moreover, the c.930+189C>T variant prompted the utilization of the pathogenic splice donor site, in both heterozygous (*Col6a1*^h/h+189T^) and homozygous (*Col6a1*^h+189T/h+189T^) tissues. Consistent with the splicing outcomes we previously described in patient samples (10), at least three mRNA species produced by the variant allele were identified (as evident in *Col6a1*^h+189T/h+189T^ mouse tissues): 1) canonical splicing (from exon 11 to 12; isoforms 2 and 3, Supplementary Figure S1C), 2) 72-nt-long pseudoexon insertion (with activation of the cryptic SA); isoforms 1 and 4, Supplementary Figure S1C), or 3) exon 11 extension (up to the pathogenic SD), for a 187-nt-long insertion predicted to be subject to nonsense-mediated decay; isoform frequency <3% and not depicted).

We next determined the percent pseudoexon inclusion in various tissues using digital PCR assays. In 8-week-old *Col6a1*^h/h+189T^ males, the percent pseudoexon inclusion varied from 7.6% to 10.6% in quadriceps, tibialis anterior, gastrocnemius and diaphragm muscles, whereas in *Col6a1*^h+189T/h+189T^ tissues, the percent pseudoexon was higher (from 15.0% to 20.8%, n=4 animals per genotype and tissue; Figure 2A). In human biopsy samples, pseudoexon expression levels were comparable to those observed in the *Col6a1*^h+189T/h+189T^ tissues (average of 24.7%, n=3; Figure 2A). Thus, the *Col6a1*^h+189T/h+189T^ genotype faithfully recapitulates the human transcript isoform expression profile. The percent pseudoexon expression in quadriceps did not significantly vary between females and males or between 8-week-old and 24-week-old mice (n=4 per sex and age group, Figure 2B).

**Figure 2.**
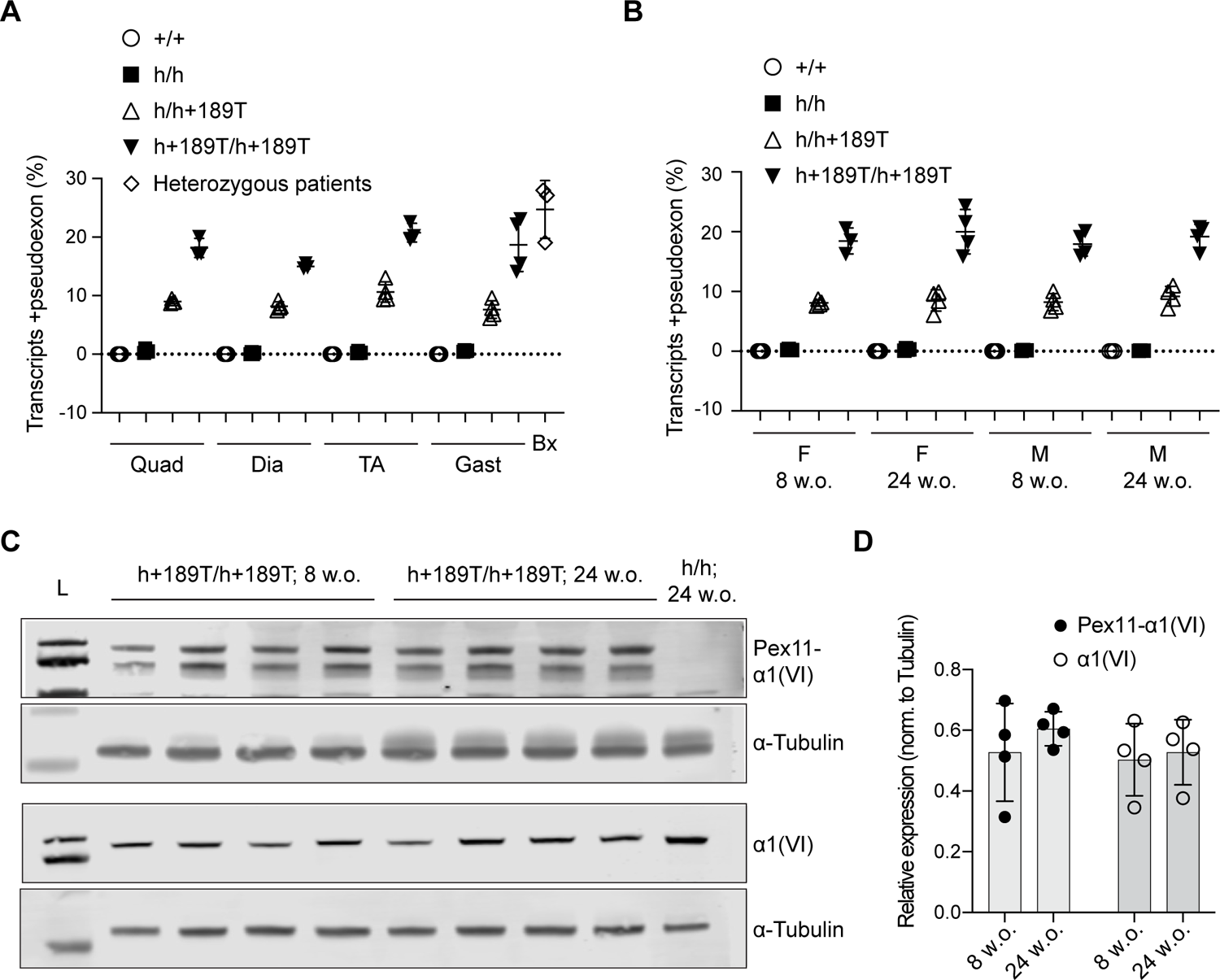
Pathogenic pseudoexon expression in the *Col6a1* h+189T model. **(A, B)** Isoform-specific digital PCR assays were used to quantify the percentage of total *Col6a1* humanized transcripts that include the 72-nt-long pseudoexon, in various tissues of 8-week-old male mice compared to heterozygous patients’ biopsies (A), and in quadricep muscles of 8-week-old and 24-week-old male and female mice (B). Data points represent individual mice (n=4 per group) or patients (n=3). Data presented as mean ± SD. Quad = quadriceps; Dia = diaphragm; TA = tibialis anterior; Gast = gastrocnemius; Bx = patient biopsies; F = females; M = males. **(C)** Translation of the pseudoexon peptide was assessed by immunoblotting with a pseudoexon-specific antibody (Pex11-α1(VI)), in quadriceps of 8-week-old and 24-week-old *Col6a1*^h+189T/h+189T^ males (n=4). **(D)** The top of the double band (in (C)) corresponds to the size of the α1(VI) protein and was used for quantification. Data points represent pseudoexon-containing α1(VI) protein expression levels normalized to tubulin (n=4 mice per group). Kolmogorov-Smirnov test was applied and showed no significant difference between the time points.

The pseudoexon encodes a 24-amino-acid-long stretch (2.7 kDa predicted) that interrupts the Gly-X-Y repeats. We previously generated an antibody (Pex11-α1(VI)) using the 24-amino-acid-long peptide as the immunogen (10, 11). The Pex11-α1(VI) antibody showed signal in quadriceps of 8-week-old and 24-week-old *Col6a1*^h+189T/h+189T^ males (top band, Figure 2C), at a molecular weight comparable to the α1(VI) chain (third panel, Figure 2C). The antibody detected an additional band of lower molecular weight that is under investigation. The levels of the pseudoexon-containing α1(VI) protein and of total collagen α1(VI) do not significantly increase from 8 to 24 weeks (Figure 2D).

### Mouse and muscle phenotyping

All *Col6a1*^h/h+189T^ or *Col6a1*^h+189T/h+189T^ mice developed normally and exhibited lifespans comparable to *Col6a1*^h/h^ and *Col6a1*^+/+^(wild-type) control animals. There were no significant body weight differences between any of the h+189T-containing genotypes and the *Col6a1*^h/h^ or *Col6a1*^+/+^ animals, from 4 weeks to 24 weeks of age, in males and females (n≥4 per group; Figure 3A, B). The muscle masses of *Col6a1*^h+189T/h+189T^ tibialis anterior and gastrocnemius muscles, however, were significantly decreased compared to *Col6a1*^+/+^ and *Col6a1*^h/h^ tissues, at 8 and 24 weeks, for both males and females (n≥4 per group; Figure 3C, D), except for males’ tibialis anterior at 24 weeks in which the muscle masses were decreased but did not reach significance. In *Col6a1*^h/h+189T^ mice, muscles weighted less compared to *Col6a1*^+/+^ and *Col6a1*^h/h^ mice, but this decrease did not consistently reach statistical significance.

**Figure 3.**
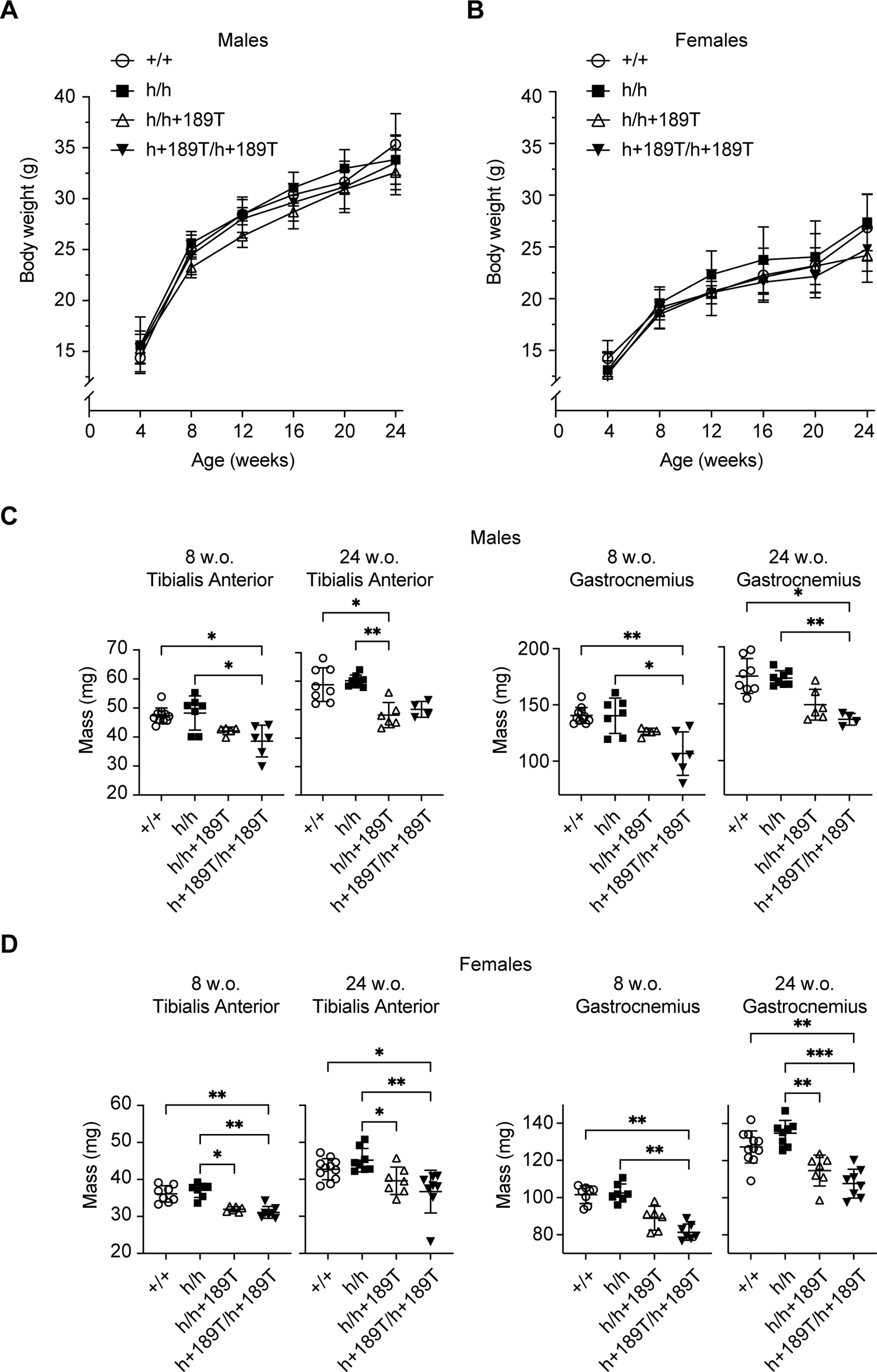
Body and muscle weights of *Col6a1* h+189T mice. **(A, B)** Whole body weights were recorded every four weeks from 4 to 24 weeks old, in males (A), and females (B). Statistical analyses were performed with one-way ANOVA followed by Tukey’s multiple comparisons test and showed no significant difference. **(C, D)** Masses of tibialis anterior and gastrocnemius muscles were recorded in 8-week-old and 24-week-old males (C) and females (D) (n=4-11). Data points represent biological replicates and data are presented as mean ± SD. Statistical analyses were performed with a Kruskal-Wallis ANOVA test followed by Dunn’s multiple comparisons. *p<0.05; **p<0.005, ***p<0.0005.

We next investigated for histopathological signs of muscular dystrophy in the humanized mice skeletal muscles using hematoxylin and eosin staining. Qualitatively, we observed mild changes in the number of centrally nucleated fibers in *Col6a1*^h/h+189T^ and in *Col6a1*^h+189T/h+189T^ mice, and we did not observe clear signs of increased fatty infiltration or increased matrix production in these genotypes (Figure 4A). The percent of centrally nucleated fibers, an indicator for degeneration/regeneration cycles that can reflect the degree of muscle pathology, was increased in *Col6a1*^h+189T/h+189T^ quadricep muscles of both males and females (n=4-5 mice per group; Figure 4B), although without reaching statistical significance.

**Figure 4.**
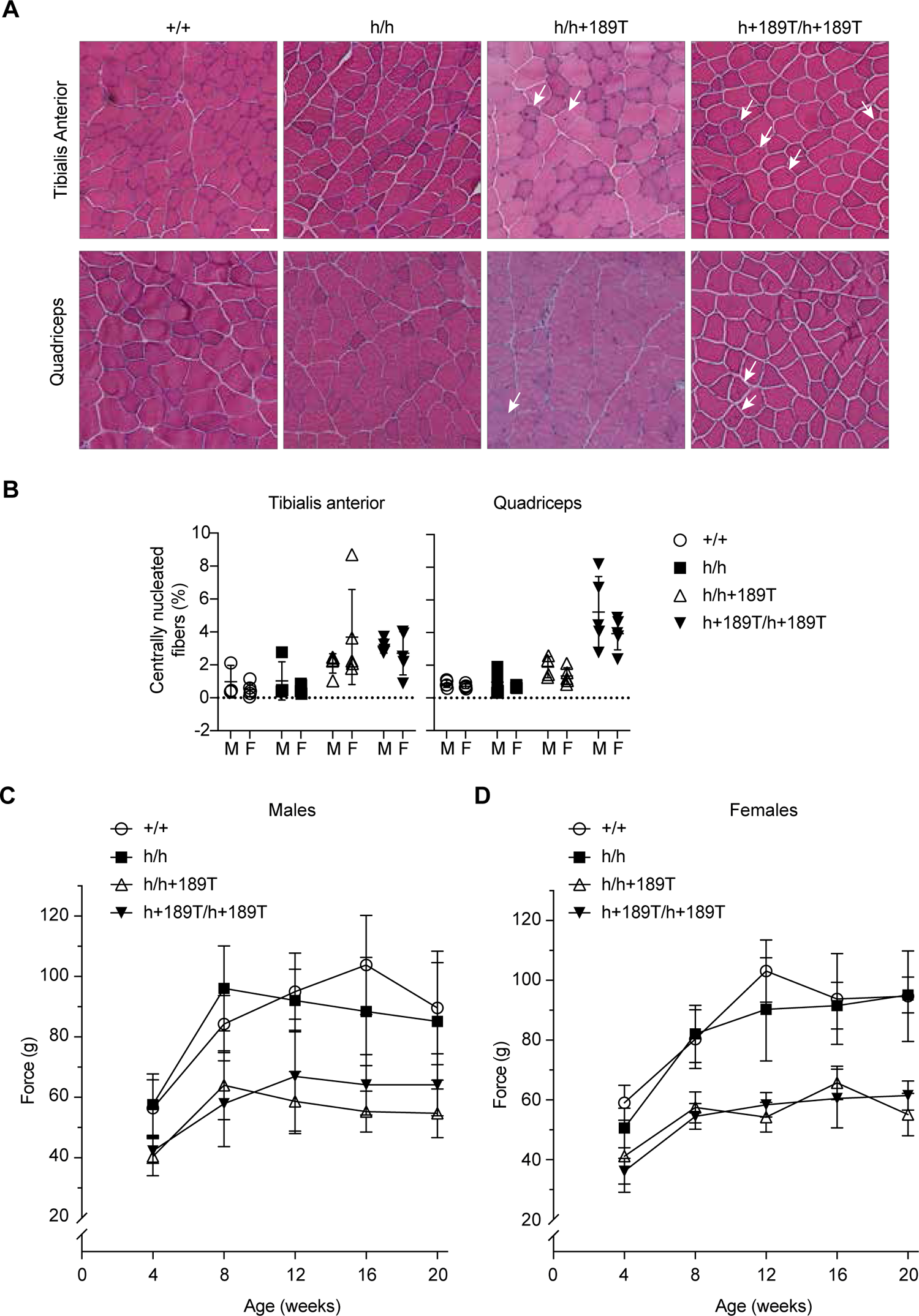
Muscle histology and function in *Col6a1* h+189T mice. **(A)** Representative images of tissue sections of tibialis anterior (top row) and quadriceps muscles (bottom row) of 24-week-old male mice stained with hematoxylin and eosin. Mild histopathological signs of muscular dystrophy were detected in *Col6a1*^h/h+189T^ and *Col6a1*^h+189T/h+189T^, evidenced by the slightly increased percent of centrally nucleated fibers (arrows). **(B)** The percent of centrally nucleated fibers was manually counted on tissue sections stained with wheat germ agglutinin and DAPI. M = males; F = females. Data points represent biological replicates (n=4-5). Data presented as mean ± SD. Statistical analyses were performed with a Kruskal-Wallis ANOVA test followed by Dunn’s multiple comparisons. **(C, D)** Absolute force of the forelimb grip strength was measured using a grip strength meter and is reported in grams (g). Grip strength was measured every 4 weeks from 4 to 24 weeks of age in males (C) and females (D). Graphs depict the mean grip strength ± SD (n=5-9).

We used the grip strength test to functionally assess forelimb muscles’ strength. Strikingly, the grip strengths of both *Col6a1*^h/h+189T^ and *Col6a1*^h+189T/h+189T^ males and females were weaker at all ages recorded between 4 and 20 weeks compared to the control groups (Figure 4C, D). The decrease in grip strength was statistically significant between the control *Col6a1*^+/+^ and *Col6a1*^h/h^ mouse groups compared to the disease *Col6a1*^h/h+189T^ and *Col6a1*^h+189T/h+189T^ mouse groups at most tested age timepoints (Supplementary Figure S2).

## Discussion

In the era of personalized medicine, genomic humanized models are becoming increasingly valuable tools for the *in vivo* testing of nucleic acid-based therapies tailored to specific human pathogenic variants (18). Modeling a pathogenic variant within a humanized gene context provides the desired target for nucleic acid-based drugs that rely on sequence recognition, such as antisense oligonucleotides or genome editing technologies. This is particularly relevant for splice-modulating therapeutics targeted at intronic sequences. In addition, humanized models enable optimizing the delivery of these human-ready drugs to the relevant tissues and cell types in the context of the disease studied.

To facilitate the preclinical development of a splice-modulating therapy for the recurrent intronic *COL6A1* variant (c.930+189C>T) as an ideal target for pseudoexon-skipping approaches, we generated a humanized mouse model that recapitulates the pseudoexon-inducing splicing event caused by the pathogenic variant. We partially humanized the *Col6a1* gene as a knock-in allele (termed “h”). Like previously generated humanized mouse models, where the canonical splice sites within a humanized gene were faithfully recognized by the mouse splicing machinery (14–16), the h allele in our model expressed full-length and functional chimeric mouse/human *Col6a1* transcripts. Modeling intronic and splice-altering variants, such as the c.930+189C>T, in humanized backgrounds can be more challenging than only inserting a humanized sequence, however (19, 20). In mice carrying a single allele of the humanized *Col6a1* that includes the intronic variant (termed “h+189T”), the pseudoexon inclusion levels did not reach the levels detected in muscle biopsies from heterozygous patients, suggesting that the pathogenic variant was underutilized in mouse tissues despite the humanization of several flanking exons and introns to maximize the likelihood of including potential *cis*-acting splicing regulators. However, when quantifying the pseudoexon inclusion levels in *Col6a1*^h+189T/h+189T^ mice tissues, we found that the pseudoexon inclusion from both alleles combined was very close to the levels detected in biopsies from the heterozygous human patients, thus providing a relevant disease model at the transcript level. In addition, we showed that in the h+189T mouse model, the pseudoexon was translated, and that the corresponding peptide was detected in proteins of molecular weight comparable to the α1(VI) protein, thus validating that the *Col6a1*^h+189T/h+189T^ mouse is a relevant preclinical model to test the on-target activity of splice-modulating therapeutics.

Collagen VI deficiency or dysfunction in various mouse models results in phenotypes milder than the clinical manifestations observed in COL6-RD patients (7, 21–24). Similarly, our *Col6a1*^h+189T/h+189T^ mice are milder in severity when compared to the Ullrich patients heterozygous for the c.930+189C>T variant (10). Nevertheless, *Col6a1*^h/h+189T^ and *Col6a1*^h+189T/h+189T^ mice show muscle weakness (assessed with the forelimb grip strength testing) and histological signs of myofiber degeneration and regeneration (seen with the moderately increased percent of centrally nucleated fibers). Grip strength assessment can be used as a longitudinal functional outcome measure when testing the effect of splice-modulating antisense oligonucleotides. It is noteworthy that the heterozygous *Col6a1*^h/h+189T^ genotype results in a phenotype not that different from the homozygous *Col6a1*^h+189T/h+189T^ genotype, even though the abundance of pseudoexon inclusion from the single allele is less. This may suggest a cumulative pathological effect of the pseudoexon peptide over time.

While this model carries the target human sequence to optimize on-target activity of nucleic acid-targeting drugs *in vivo*, it is of limited use to assess sequence-dependent off-target effects, since the genomic background consists of the mouse genome. Nevertheless, sequence-dependent off-targets can be predicted and assessed in cellular models. Moreover, toxicity due to components other than the primary sequence (based on the chemistry, for example) can be tested in organisms such as non-human primates.

In the *COL6A1* gene, dominant-negative variants occur within the N-TH, mostly between exons 9 and 14. Other than the intron 11 variant described here, common *COL6A1* variants also located in this N-TH domain include splice-site variants that result in exon 14 skipping (25, 26), and the G284R and G293R substitutions in exons 9 and 10, respectively (27, 28). Notably, since the three *COL6* genes are haplosufficient (1, 2), *COL6* variants like the ones described above are also promising targets for nucleic acid therapeutics, in particular for therapeutics that mediate allele-specific silencing, such as small interfering RNAs (29, 30), RNase H-recruiting gapmers (31), and gene editing (32). Using as a template the humanized knock-in allele we created (‘h’), it is now feasible to generate humanized models for each one of these variants on this background to then test human-ready antisense therapies. We anticipate that our unique humanized knock-in model will accelerate the development of nucleic acid therapeutics for COL6-RDs.

## Material and Methods

### Generation of the *Col6a1*^h/h^, *Col6a1*^h/h+189T^ and *Col6a1*^h+189T/h+189T^ mice

The mouse and human sequences were aligned using NCBI Blastn suite. All mouse procedures were approved by the NINDS Animal Care and Use Committee (Animal Safety Protocol #1556). Manipulations of recombinant DNA was approved by the National Institutes of Health Institutional Biosafety Committee (Registration #RD-17-VI-01). The humanized knock-in mice were created using homology recombination induced by CRISPR/Cas9 at the National Human Genome Research Institute (NHGRI) Transgenic and Gene Editing Core facility. A combination of two guide RNAs were prepared with SpCas9 into ribonucleoprotein complexes and delivered to C57BL6/N x C57BL6/J zygotes by pronuclear injections, together with a targeted construct. The guide RNAs were selected to cleave 117 bp upstream of exon 9 and 66 bp downstream of exon 14 of the mouse *Col6a1* and were synthesized by Horizon (Horizon Discovery/Dharmacon, Lafayette, CO). We created two targeted constructs containing the human genomic sequence from intron 8 to intron 14 of the *COL6A1* gene flanked by 800 bp of mouse homology arms. One construct contained the reference cytosine at position c.930+189 (here referred to as “h”), and the other construct carried the pathogenic thymine at the position (here referred to as “h+189T”). The constructs included 160 bp of human intron 8 sequence upstream of exon 9 (preserving two potential branch point sites) and 65 bp of sequence downstream of exon 14. Each of the two targeted constructs was 3,520 bp in length and were synthesized and cloned (Epoch LifeScience, Missouri City, TX). The plasmids were prepared and purified by 2X CsCl gradients (Lofstrand Labs, Gaithersburg, MD). The targeted vectors were isolated by restriction enzymes prior to pronuclear injection. Zygotes were implanted into CBy6F1 pseudo-pregnant females. F0 pups DNA was extracted from tail biopsies and was analyzed by PCR and Sanger sequencing. For each of the two constructs, a single pup was identified with the expected sequence, and was bred with a C57BL/6J animal. Animals were backcrossed for 5 generations with C57BL/6J animals. Breeding colonies were maintained as *Col6a1*^h/h^ x *Col6a1*^h/h^; *Col6a1*^+/h+189T^ x *Col6a1*^+/h+189T^; or *Col6a1*^h/h^ x *Col6a1*^+/h+189T^ to generate all genotypes used in the study (*Col6a1*^+/+^, *Col6a1*^h/h^, *Col6a1*^h/h+189T^, *Col6a1*^h+189T/h+189T^). Animals were housed at the Intramural NINDS Building 35 vivarium under the following conditions: 12-hour light/dark cycles, 20-25 °C room temperature, 40-65% relative humidity, access to untreated drinking water and chow *ad libitum*. Genotyping was performed with a PCR amplification combining a single forward primer located in the mouse intron 8 upstream of the knock-in region (5’-cagttcagccttgatgcaaa-3’) with two reverse primers each hybridizing specifically a different allele, either the mouse wild-type allele (5’-gcagaggaaatcagctcagg-3’) or the humanized knock-in allele (5’-taaagcaccttccccatcac-3’). The difference in molecular weight of these two amplicons (66 bp) was sufficient to separate and visualize with a 2.5% agarose gel electrophoresis and to identify three genotypes: unmodified wild-type, and either heterozygous or homozygous for the humanized knock-in allele. When needed, we confirmed *Col6a1*^h/h^, *Col6a1*^h/h+189T^ and *Col6a1*^h+189T/h+189T^ mouse genotypes using a custom genotyping assay (ThermoFisher, Waltham, MA) in which the probes detecting the ‘T’ (h+189T) or the ‘C’ (h) alleles were labeled with different fluorescent dyes (FAM and VIC, respectively). These PCR products were detected at end point on the QuantStudio6 Real-Time PCR instrument (ThermoFisher).

### Mouse phenotyping

The mice body weights and forelimb grip strength measurements were assessed on the same day once a month (every 28 days). The mice were weighed before the grip strength assessment. The mice forelimb grip strength was measured with a digital grip strength meter (Bio-GS3, Bioseb, Pinellas Park, FL) using a standard grip strength assay (33, 34). The middle three values of five measurements were averaged and reported as the absolute force generated in grams (g). One-way ANOVA test was used followed by Tukey’s HSD as a post-hoc test to determine statistical significance for body weights and grip strengths values.

### Patients

Specimen collection and use was approved by the National Institutes of Health Institutional Review Board (Protocol #12-N-0095).

### Tissue processing

For human muscle biopsies, RNA was isolated from fast-frozen tissue sections (prepared on the CM1950 cryostat, Leica, Wetzlar, Germany) using Trizol (ThermoFisher, Waltham, MA). For mouse specimen, muscles were either fast-frozen in dry ice-prechilled isopentane (for histology processing) or cut into ∼20 mg pieces and snap-frozen in liquid nitrogen (for RNA and protein processing). To isolate RNA, tissues were homogenized in Trizol using the Red Eppendorf Lysis kits and the Bullet Blender (Next Advanced, Troy, NY). RNA was then isolated using manufacturer guidelines. To isolate proteins, tissues were homogenized and lysed in a Urea/Thiourea Buffer (7 M urea, 2 M thiourea, 4% (w/v) CHAPS in 30 mM Tris-HCl, pH 8.5) (35) containing 1X cOmplete protease inhibitor cocktail (MilliporeSigma, Burlington, MA), using the Green Eppendorft Lysis kits and the Bullet Blender (Next Advance). Reagent DX (Qiagen, Germantown, MD) was added to the lysis buffer at 0.5% v/v to prevent foaming during homogenization.

### Long-range sequencing

A panel of 101 120-mer probes was designed to span the entire coding sequence of the mouse/human hybrid *Col6a1* coding sequence. This set also included probes specific to each allele at the c.930+189 position (IDT xGen Pool Design, Integrated DNA Technologies, Coralville, IA). RNA samples were first cleaned up using bead purification (AMPure RNA-XP, Beckman Coulter, Indianapolis, IN). First-strand synthesis, transcripts capture, library preparation and PacBio Sequel II sequencing were performed at Sequencing Facility – Long Read Technology Center for Cancer Research of the National Cancer Institute (Frederick, MD). Reads were aligned to a custom reference genome. A publicly available workflow (v4.0) to identify transcripts in PacBio single-molecule sequencing data was used (https://github.com/PacificBiosciences/IsoSeq). The specific workflow used included two main execution steps, Clustering (https://isoseq.how/clustering/cli-workflow.html) and Classification (https://isoseq.how/classification/workflow.html). Tools used as part of this workflow included those supported in the SMRT® Analysis software suite v13.0.0.207600 (https://www.pacb.com/support/software-downloads/). All computational analyses were performed on the NIH HPC Biowulf cluster (http://hpc.nih.gov). Sashimi plots were generated using ggsashimi (36). BAM files were loaded into VIsoQLR (37), which was used to detect and quantify splice isoform species using standard parameters, except for adjustments of the base-pair lengths made to the detected terminal exons to compensate for sequencing biases.

### Digital PCRs (dPCRs)

RNA was converted to complementary DNA (cDNA) using SuperScript IV Reverse Transcriptase (ThermoFisher). Using PrimerQuest Tool (Integrated DNA Technologies) to design custom taqman-based assays, we selected two assays that specifically amplify either pseudoexon-containing transcripts only, or any humanized transcript (spanning exons 12 to 14), ordered as PrimeTime assays. We purchased a pre-designed mouse *Csnk2a2* assay (Mm.PT.58.10226157) as the housekeeping control (Integrated DNA Technologies). We performed quantitative PCRs on complementary DNA (cDNA) samples by combining each isoform-specific assay with the housekeeping assay. Reaction partitioning and amplification, and fluorescence detection were done using the QIAcuity digital PCR instrument (Qiagen). For each assay, thresholds were determined manually and applied to all samples. The concentration values (copies/µL) obtained for each isoform-specific assay were normalized to the concentration obtained for the housekeeping assay. Percent pseudoexon inclusion was calculated by dividing the pseudoexon-containing transcript concentration by the concentration of all humanized transcripts. For the human biopsy samples, the same isoform-specific assays were utilized, whereas a human *CSNK2A2* pre-designed assay (Hs00176505_m1, ThermoFisher) was used as the housekeeping control.

### Immunoblots

Protein lysates were quantified using the 660 nm Protein Assay Kit (Pierce/ThermoFisher) and read on the Victor Nivo multimode plate reader (PerkinElmer, Waltham, MA). Protein samples were diluted in a loading dye containing 100 mM dithiothreitol (DTT). Samples were boiled at 95°C for 5 min before loading (60 µg) on 4-12% Bis-Tris gel (ThermoFisher) and ran with 1X MES buffer (ThermoFisher) for 45 min at 125 volts. Transfer to PVDF membrane (MilliporeSigma) was done on the XCell II Blot module (ThermoFisher) in 1X Transfer buffer (ThermoFisher) supplemented with 5% methanol at 25 volts for 90 min. Membranes were blocked in Intercept Blocking Buffer (LI-COR, Lincoln, NE) for 1 hr before adding primary antibodies: anti-tubulin (T5168 (MillporeSigma), diluted 1:10,000 in blocking solution) or anti-Collagen VI (ab199720 (Abcam, Cambridge, UK), diluted 1:1,000 in blocking solution) for 90 min. The affinity-purified anti-rabbit pseudoexon antibody (Pex11-a1(VI)), gift from Raimund Wagener, was diluted 1:1,000 in blocking solution. Secondary antibodies used were 680RD goat anti-rabbit IgG (926-68071, LI-COR), diluted 1:10,000 or 800CW goat anti-mouse IgG (926-32210, LI-COR) diluted 1:15,000 in blocking solution and incubated for 1 hr. Washes were done in phosphate-buffered saline containing 0.05% tween-20 (MP Biomedicals, Santa Ana, CA). Detection was done on the Odyssee Clx Imaging System (LI-COR) using the Image Studio Software Ver 5.2 (LI-COR). Areas under the curves were measured with Fiji and used for quantification. Values for collagen α1(VI) were normalized to tubulin. Unpaired Kolmogorov-Smirnov test was applied.

### Histology

Tissues were sectioned (10 µm) using the CM1950 cryostat (Leica). For hematoxilin and eosin staining, frozen slides were first thawed and incubated in filtered Harris Hematoxylin (VWR, Radnor, PA) for 8 min before being briefly dipped in water containing 0.2% ammonium hydroxide for 50 sec. The samples were then put through a series of additional dips: 95% EtOH for 5 sec, Eosin (VWR) for 1 min, 95% EtOH for 5 sec, followed by 95% EtOH for 2 min (3X), 100% EtOH for 2 min (3X), and Xylene for 2 min (3X). Specimen were covered with Permount (Fisher Scientific, Hampton, NH). For the quantification of centrally nucleated fibers, a wheat germ agglutinin and DAPI stain was used. Briefly, slides were fixed with 4% PFA (Electron Microscopy Sciences, Hatfield, PA) for 10 min and blocked with phosphate-buffered saline solution containing 5% normal goat serum and 0.3% Triton X-100 (Sigma-Aldrich, St. Louis, MO) for 30 min. Secondary antibodies used were WGA-488 (ThermoFisher) diluted 1:500 and DAPI in blocking buffer and incubated with samples for 15 min. Slides were mounted with Fluoromount-G (Southern Biotech, Birmingham, AL). The percent of centrally nucleated fibers was manually counted with the help of the Fiji software. Kruskal-Wallis test followed by Dunn’s multiple comparisons was applied.

### Microscopy

Images were captured using the 20X objective on the Eclipse Ti microscope system (Nikon, Tokyo, Japan) and using the NIS-Elements AR Software (Nikon).

## Acknowledgements

We thank Kory Johnson for the bioinformatics support and Gina Norato for the biostatistics consultation and support. We thank Raimund Wagener for providing the Pex11-a1(VI) antibody. This work was supported through a Conditional Gift from Muscular Dystrophy UK (Project Reference 16CollVI-PG24-0123) to C.G.B. This research was made possible through the NIH Medical Research Scholars Program, a public-private partnership supported jointly by the NIH and contributions to the Foundation for the NIH from the Doris Duke Charitable Foundation, Genentech, the American Association for Dental Research, the Colgate-Palmolive Company, and other private donors. This work was supported by the Division of Intramural Research of the NIH, NINDS (1ZIANS003129). The content is solely the responsibility of the author(s) and does not necessarily represent the official views of the National Institutes of Health.

## References

1. Bonnemann CG. The collagen VI-related myopathies: muscle meets its matrix. Nature reviews Neurology. 2011;7(7):379–90.

2. Mohassel P, Foley AR, Bonnemann CG. Extracellular matrix-driven congenital muscular dystrophies. Matrix biology: journal of the International Society for Matrix Biology. 2018;71–72:188-204.

3. Allamand V, Brinas L, Richard P, Stojkovic T, Quijano-Roy S, Bonne G. ColVI myopathies: where do we stand, where do we go? Skeletal muscle. 2011;1:30.

4. Zou Y, Zhang RZ, Sabatelli P, Chu ML, Bonnemann CG. Muscle interstitial fibroblasts are the main source of collagen VI synthesis in skeletal muscle: implications for congenital muscular dystrophy types Ullrich and Bethlem. Journal of neuropathology and experimental neurology. 2008;67(2):144–54.

5. Braghetta P, Ferrari A, Fabbro C, Bizzotto D, Volpin D, Bonaldo P, et al. An enhancer required for transcription of the Col6a1 gene in muscle connective tissue is induced by signals released from muscle cells. Experimental cell research. 2008;314(19):3508–18.

6. Uapinyoying P, Hogarth M, Battacharya S, Mazala DAG, Panchapakesan K, Bonnemann CG, et al. Single-cell transcriptomic analysis of the identity and function of fibro/adipogenic progenitors in healthy and dystrophic muscle. iScience. 2023;26(8):107479.

7. Lamande SR, Bateman JF. Collagen VI disorders: Insights on form and function in the extracellular matrix and beyond. Matrix biology: journal of the International Society for Matrix Biology. 2018;71–72:348-67.

8. Cescon M, Gattazzo F, Chen P, Bonaldo P. Collagen VI at a glance. Journal of cell science. 2015;128(19):3525–31.

9. Cummings BB, Marshall JL, Tukiainen T, Lek M, Donkervoort S, Foley AR, et al. Improving genetic diagnosis in Mendelian disease with transcriptome sequencing. Science translational medicine. 2017;9(386).

10. Bolduc V, Foley AR, Solomon-Degefa H, Sarathy A, Donkervoort S, Hu Y, et al. A recurrent COL6A1 pseudoexon insertion causes muscular dystrophy and is effectively targeted by splice-correction therapies. JCI Insight. 2019;4(6).

11. Freiburg CD, Solomon-Degefa H, Freiburg P, Mörgelin M, Bolduc V, Schmitz S, et al. The UCMD-Causing COL6A1 (c.930+189C>T) Intron Mutation Leads to the Secretion and Aggregation of Single Mutated Collagen VI 1 Chains. Human mutation. 2023;2023.

12. Zhu F, Nair RR, Fisher EMC, Cunningham TJ. Humanising the mouse genome piece by piece. Nature communications. 2019;10(1):1845.

13. Devoy A, Bunton-Stasyshyn RK, Tybulewicz VL, Smith AJ, Fisher EM. Genomically humanized mice: technologies and promises. Nature reviews Genetics. 2011;13(1):14–20.

14. tHoen PA, de Meijer EJ, Boer JM, Vossen RH, Turk R, Maatman RG, et al. Generation and characterization of transgenic mice with the full-length human DMD gene. The Journal of biological chemistry. 2008;283(9):5899–907.

15. Devoy A, Price G, De Giorgio F, Bunton-Stasyshyn R, Thompson D, Gasco S, et al. Generation and analysis of innovative genomically humanized knockin SOD1, TARDBP (TDP-43), and FUS mouse models. iScience. 2021;24(12):103463.

16. Luo JL, Yang Q, Tong WM, Hergenhahn M, Wang ZQ, Hollstein M. Knock-in mice with a chimeric human/murine p53 gene develop normally and show wild-type p53 responses to DNA damaging agents: a new biomedical research tool. Oncogene. 2001;20(3):320–8.

17. Veltrop M, van Vliet L, Hulsker M, Claassens J, Brouwers C, Breukel C, et al. A dystrophic Duchenne mouse model for testing human antisense oligonucleotides. PloS one. 2018;13(2):e0193289.

18. Aartsma-Rus A, van Putten M. The use of genetically humanized animal models for personalized medicine approaches. Disease models & mechanisms. 2019;13(2).

19. Slijkerman R, Goloborodko A, Broekman S, de Vrieze E, Hetterschijt L, Peters T, et al. Poor Splice-Site Recognition in a Humanized Zebrafish Knockin Model for the Recurrent Deep-Intronic c.7595-2144A>G Mutation in USH2A. Zebrafish. 2018;15(6):597–609.

20. Garanto A, van Beersum SE, Peters TA, Roepman R, Cremers FP, Collin RW. Unexpected CEP290 mRNA splicing in a humanized knock-in mouse model for Leber congenital amaurosis. PloS one. 2013;8(11):e79369.

21. Bonaldo P, Braghetta P, Zanetti M, Piccolo S, Volpin D, Bressan GM. Collagen VI deficiency induces early onset myopathy in the mouse: an animal model for Bethlem myopathy. Human molecular genetics. 1998;7(13):2135–40.

22. Pan TC, Zhang RZ, Arita M, Bogdanovich S, Adams SM, Gara SK, et al. A mouse model for dominant collagen VI disorders: heterozygous deletion of Col6a3 Exon 16. The Journal of biological chemistry. 2014;289(15):10293–307.

23. Mohassel P, Rooney JE, Zou Y, Johnson K, Norato G, Hearn H, et al. Collagen type VI regulates TGFβ bioavailability in skeletal muscle. bioRXiv. 2023.

24. Noguchi S, Ogawa M, Malicdan MC, Nonaka I, Nishino I. Muscle Weakness and Fibrosis Due to Cell Autonomous and Non-cell Autonomous Events in Collagen VI Deficient Congenital Muscular Dystrophy. EBioMedicine. 2017;15:193–202.

25. Pepe G, Giusti B, Bertini E, Brunelli T, Saitta B, Comeglio P, et al. A heterozygous splice site mutation in COL6A1 leading to an in-frame deletion of the alpha1(VI) collagen chain in an italian family affected by bethlem myopathy. Biochemical and biophysical research communications. 1999;258(3):802–7.

26. Lampe AK, Bushby KM. Collagen VI related muscle disorders. Journal of medical genetics. 2005;42(9):673–85.

27. Lampe AK, Dunn DM, von Niederhausern AC, Hamil C, Aoyagi A, Laval SH, et al. Automated genomic sequence analysis of the three collagen VI genes: applications to Ullrich congenital muscular dystrophy and Bethlem myopathy. Journal of medical genetics. 2005;42(2):108–20.

28. Butterfield RJ, Foley AR, Dastgir J, Asman S, Dunn DM, Zou Y, et al. Position of Glycine Substitutions in the Triple Helix of COL6A1, COL6A2, and COL6A3 is Correlated with Severity and Mode of Inheritance in Collagen VI Myopathies. Human mutation. 2013;34(11):1558–67.

29. Noguchi S, Ogawa M, Kawahara G, Malicdan MC, Nishino I. Allele-specific Gene Silencing of Mutant mRNA Restores Cellular Function in Ullrich Congenital Muscular Dystrophy Fibroblasts. Molecular therapy Nucleic acids. 2014;3:e171.

30. Bolduc V, Zou Y, Ko D, Bonnemann CG. siRNA-mediated Allele-specific Silencing of a COL6A3 Mutation in a Cellular Model of Dominant Ullrich Muscular Dystrophy. Molecular therapy Nucleic acids. 2014;3:e147.

31. Marrosu E, Ala P, Muntoni F, Zhou H. Gapmer Antisense Oligonucleotides Suppress the Mutant Allele of COL6A3 and Restore Functional Protein in Ullrich Muscular Dystrophy. Molecular therapy Nucleic acids. 2017;8:416–27.

32. Lopez-Marquez A, Morin M, Fernandez-Penalver S, Badosa C, Hernandez-Delgado A, Natera-de Benito D, et al. CRISPR/Cas9-Mediated Allele-Specific Disruption of a Dominant COL6A1 Pathogenic Variant Improves Collagen VI Network in Patient Fibroblasts. Int J Mol Sci. 2022;23(8).

33. Aartsma-Rus A, van Putten M. Assessing functional performance in the mdx mouse model. J Vis Exp. 2014(85).

34. Castro B, Kuang S. Evaluation of Muscle Performance in Mice by Treadmill Exhaustion Test and Whole-limb Grip Strength Assay. Bio Protoc. 2017;7(8).

35. Miskiewicz EI, MacPhee DJ. Lysis Buffer Choices Are Key Considerations to Ensure Effective Sample Solubilization for Protein Electrophoresis. Methods in molecular biology. 2019;1855:61–72.

36. Garrido-Martin D, Palumbo E, Guigo R, Breschi A. ggsashimi: Sashimi plot revised for browser- and annotation-independent splicing visualization. PLoS computational biology. 2018;14(8):e1006360.

37. Nunez-Moreno G, Tamayo A, Ruiz-Sanchez C, Corton M, Minguez P. VIsoQLR: an interactive tool for the detection, quantification and fine-tuning of isoforms in selected genes using long-read sequencing. Human genetics. 2023;142(4):495–506.

