## Supplementary Figures for "A humanized knock-in *Col6a1* mouse recapitulates a deep-intronic splice-activating variant"

**Supplementary Figure S1. Isoforms detection in quadriceps using long-read RNA sequencing.**

**(A)** Custom builds for the *Mus musculus* (mm)/*Homo sapiens* (hs) *Col6a1* transcripts. The bottom transcript includes the 72-nt-long pseudoexon (PE). **(B-C)** Transcript isoforms in quadriceps from an 8-week-old *Col6a1*$^{h/h}$ mouse (B) and an 8-week-old *Col6a1*$^{h+189T/h+189T}$ mouse (C) were automatically detected by VIsoQLR from the BAM files. Values represent the percent of classified reads. Only isoforms with frequency greater than 3% are depicted.

**Supplementary Figure S2. Grip strength of *Col6a1* h+189T mice.**

Absolute forelimb grip strength force of 4-, 8-, and 20-week-old male (A) or female (B) mice. Each data point represents one individual mouse. Data are presented as mean ± SD (n=4-8 mice). Statistical analyses were performed with one-way ANOVA followed by Tukey's multiple comparisons test. *p<0.05; **p<0.005, ***p<0.0005; ****p<0.0001.

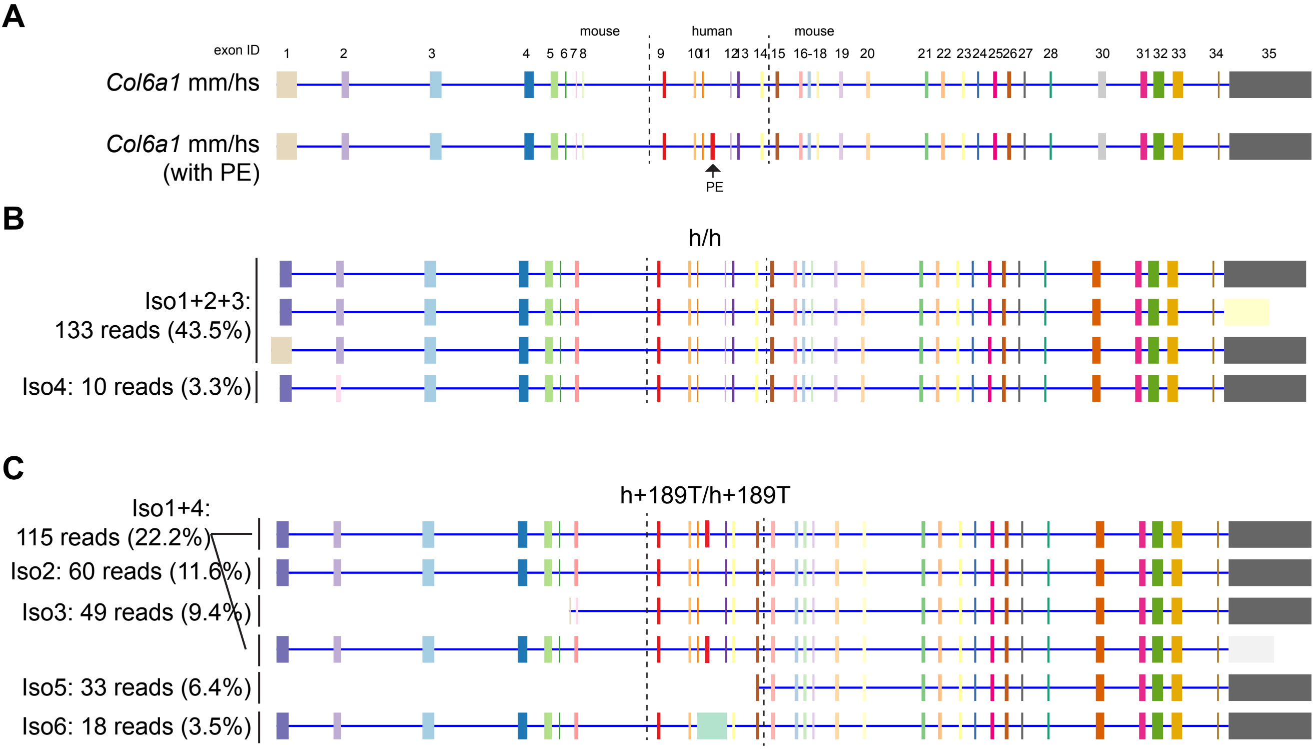


### Supplementary Figure S1

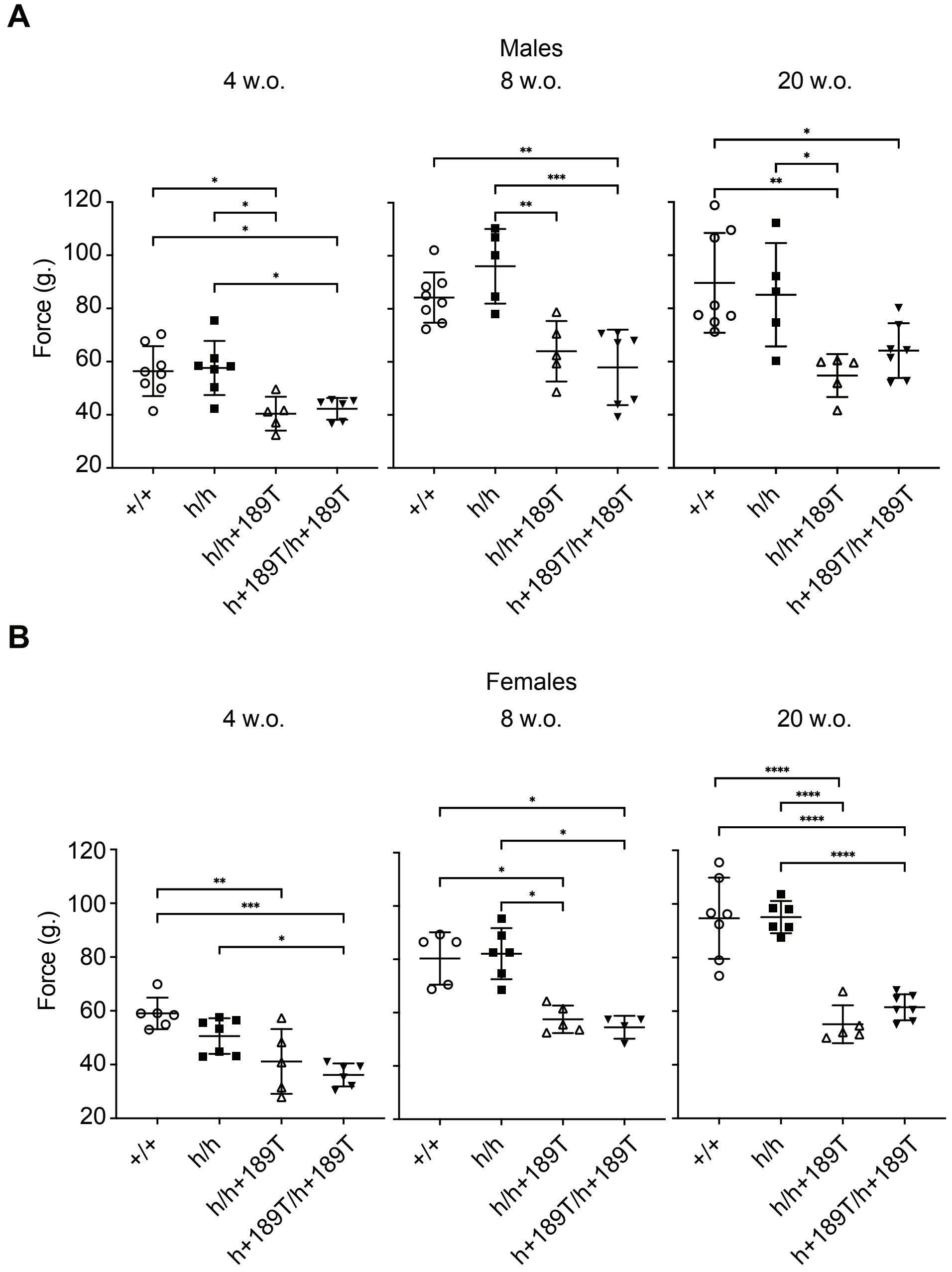


**Supplementay Figure S2**
